# Innate visual preferences and behavioral flexibility in *Drosophila*

**DOI:** 10.1101/336412

**Authors:** Martyna J. Grabowska, James Steeves, Julius Alpay, Matthew Van De Poll, Deniz Ertekin, Bruno van Swinderen

## Abstract

Visual decision-making in animals is influenced by innate preferences as well as experience. Interaction between hard-wired responses and changing motivational states determines whether a visual stimulus is attractive, aversive, or neutral. It is however difficult to separate the relative contribution of nature versus nurture in experimental paradigms, especially for more complex visual parameters such as the shape of objects. We used a closed-loop virtual reality paradigm for walking *Drosophila* flies to uncover innate visual preferences for the shape and size of objects, in a recursive choice scenario allowing the flies to reveal their visual preferences over time. We found that *Drosophila* flies display a robust attraction / repulsion profile for a range of objects sizes in this paradigm, and that this visual preference profile remains evident under a variety of conditions and persists into old age. We also demonstrate a level of flexibility in this behavior: innate repulsion to certain objects could be transiently overridden if these were novel, although this effect was only evident in younger flies. Finally, we show that a reward circuit in the fly brain, *Drosophila* neuropeptide F (dNPF), can be recruited to guide visual decision-making. Optogenetic activation of dNPF-expressing neurons converted a visually repulsive object into a more attractive object. This suggests that dNPF activity in the *Drosophila* brain guides ongoing visual choices, to override innate preferences and thereby provide a necessary level of behavioral flexibility in visual decision-making.

## Introduction

Animals continuously make decisions to survive in a dynamic environment, to for example successfully locate an adequate food source, find a way home, or avoid something dangerous. Behavioral choices are guided by innate preferences or ‘instinct’, as well as by more flexible cognitive processes such as attention (VanRullen and Thorpe, 2001; Smith and Ratcliff, 2009), learning, and memory (Tobler *et al.*, 2006; Euston, Gruber and McNaughton, 2012; O’Doherty, Cockburn and Pauli, 2017; Odoemene, Nguyen and Churchland, 2017). Instinct and experience together determine the valence of stimuli and therefore assign negative or positive associations (Lee, McGreevy and Barraclough, 2005; Gold and Shadlen, 2007; Xie and Padoa-Schioppa, 2016). Typically, negative and positive associations to stimuli result in opposing behavioral actions: animals move towards attractive stimuli and away from aversive stimuli. In animal learning experiments, these rudimentary behaviors are usually tested by using a Pavlovian conditioning paradigm, whereby one of two ‘neutral’ stimuli are provided a valence cue (a punishment or reward) in order to demonstrate increased attraction (or repulsion) towards that stimulus, compared to the other (Tully and Quinn, 1985; Balleine and Dickinson, 1998; Dickinson and Balleine, 2002; Rangel, Camerer and Montague, 2008). However, the valence of stimuli is not necessarily hard-wired (Janak and Tye, 2015). Inherently attractive objects can become less attractive over time due to habituation, or can become repulsive if associated with punishment. Similarly, inherently repulsive objects might become transiently worth paying attention to (Redondo *et al.*, 2014). Such flexibility seems to be a feature of all animal brains, to allow for adaptive decision-making based on experience.

Recent studies suggest that circuits in the central brain of insects, in the central complex (CC), are involved in decision making (Guo *et al.*, 2016; Sun *et al.*, 2017). These insect circuits display some functional similarities to the mammalian basal ganglia (Stephenson-Jones *et al.*, 2011; Strausfeld and Hirth, 2013; Anderson, Laurent and Yantis, 2014; Barron *et al.*, 2015), especially with regard to the regulation of valence-based decision making (VanRullen and Thorpe, 2001; Gold and Shadlen, 2007; Rangel, Camerer and Montague, 2008; Dan *et al.*, 2011; Guitart-Masip *et al.*, 2012; O’Doherty, Cockburn and Pauli, 2017). Also like the basal ganglia, the insect CC is involved in multisensory integration; it is understood to be involved in sleep (Nitz *et al.*, 2002; Donlea *et al.*, 2011, 2018), learning and memory (Liu *et al.*, 2006; van Swinderen, 2007; Krashes *et al.*, 2009; Zhang *et al.*, 2013; Weir, Schnell and Dickinson, 2014; Rohwedder *et al.*, 2015), navigation and orientation (Seelig and Jayaraman, 2015), and action-selection (Gurney, Prescott and Redgrave, 2001; Gerfen and Surmeier, 2011; Barron *et al.*, 2015; Barron and Klein, 2016). These diverse functions are regulated in the brain for both mammals and insects by monoamines, (Keene and Waddell, 2007; Waddell, 2010; Kahsai and Winther, 2011; Weir, Schnell and Dickinson, 2014; Ichinose *et al.*, 2015; Gøtzsche and Woldbye, 2016), gamma-Aminobutyric acid (GABA)(Gurney, Prescott and Redgrave, 2001; Root *et al.*, 2008; Zhang *et al.*, 2013; Guo *et al.*, 2016) and neuropeptides (Bannon *et al.*, 2000; Krashes *et al.*, 2009; Gøtzsche and Woldbye, 2016; Chung *et al.*, 2017; Shao *et al.*, 2017). Whether these neuromodulators and neuropeptides play a direct role in controlling valence based decision-making in the insect CC remains unclear.

*Drosophila* neuropeptide F (dNPF) is the homologue of mammalian neuropeptide Y (NPY) (Garczynski *et al.*, 2002), which is involved in signaling food satiety levels as well as regulation of fear and anxiety (Redrobe, Dumont and Quirion, 2002; Primeaux *et al.*, 2005; Gøtzsche and Woldbye, 2016). In *Drosophila*, an increase in dNPF levels in the brain has been associated with increased aggression (Dierick and Greenspan, 2007), arousal (Chung *et al.*, 2017), and reward learning (Krashes *et al.*, 2009; Shao *et al.*, 2017). Further, dNPF modulates olfactory learning by inhibiting DA neurons that provide positive and negative valence cues to the mushroom bodies (MB) (Zhang *et al.*, 2007; Krashes *et al.*, 2009; Hattori *et al.*, 2017), a structure that has primarily been associated with olfactory memory (Keene and Waddell, 2007). Interestingly, dNPF-expressing neurons also project to the fan-shaped body (FB) (Krashes *et al.*, 2009; Kahsai and Winther, 2011), a CC neuropil associated with arousal (Donlea *et al.*, 2011; Liu *et al.*, 2012) as well as visual behavior (Liu *et al.*, 2006; Weir, Schnell and Dickinson, 2014). This suggests that dNPF might provide valence cues for visual stimuli, in order to guide visual decision-making.

In this study, we investigate visual preferences in *Drosophila*, to study flexibility in valence-based choice behavior along one visual stimulus parameter: object height. Using a closed-loop virtual reality arena for tethered, walking flies, we find that flies display robust attraction or repulsion behaviors to very specific object heights. We then show that these apparently hard-wired visual preferences can be modified by experience and controlled or overwritten by optogenetic activation of dNPF-neurons.

## Results

### Visual fixation in a closed-loop virtual reality environment for walking flies

A female *Drosophila melanogaster* fly was positioned on an air-supported ball in the center of a hexagonal LED arena (Fig. 1A, B). The fly was presented with a visual stimulus (a dark bar on a lit green background, 15°wide and 60° height). Walking of the fly resulted in forward, lateral, and turning movements of the ball. These movements were translated into corresponding movements of the visual stimulus displayed by the LED arena via a camera-based closed-loop interface (FicTrac (Moore et al., 2014), Fig. 1C). This setup allowed the fly to keep the visual stimulus in the frontal visual field (FVF) voluntarily, so we could assess fixation and attention-like parameters. Flies rapidly learned to fixate on the virtual object and increased their fixation significantly over three consecutive 2-minute trials (Fig. 1D, E). To ensure that flies actively fixated on the object, we introduced visual perturbations, where the stimulus was randomly moved by 60° to the left or to the right every 10-30 seconds. If the flies were actively attending to the stimulus, they rapidly returned it to their FVF within 10s (Fig. 1F, and see Methods). We found no significant difference in the proportion of successful returns after the perturbations (Mean±SD: Trial1: 83.5±20.6, Trial2: 79.9±28.7, Trial3: 86.8±24.4, ANOVA) or the time taken to return the stimulus to the FVF (Medians: Trial1; 5.8s, Trial2: 6.1s, Trial3: 6.0s, Kruskal-Wallis test), between the three trials (Fig. 1G, H). Further, there was no significant difference in walking speed during the three trials (Medians: Trial1: 2.7mm/s, Trial2: 2.6mm/s, Trial3: 2.7mm/s, Kruskal-Wallis Test) (Fig. 1I).

**Figure 1.**
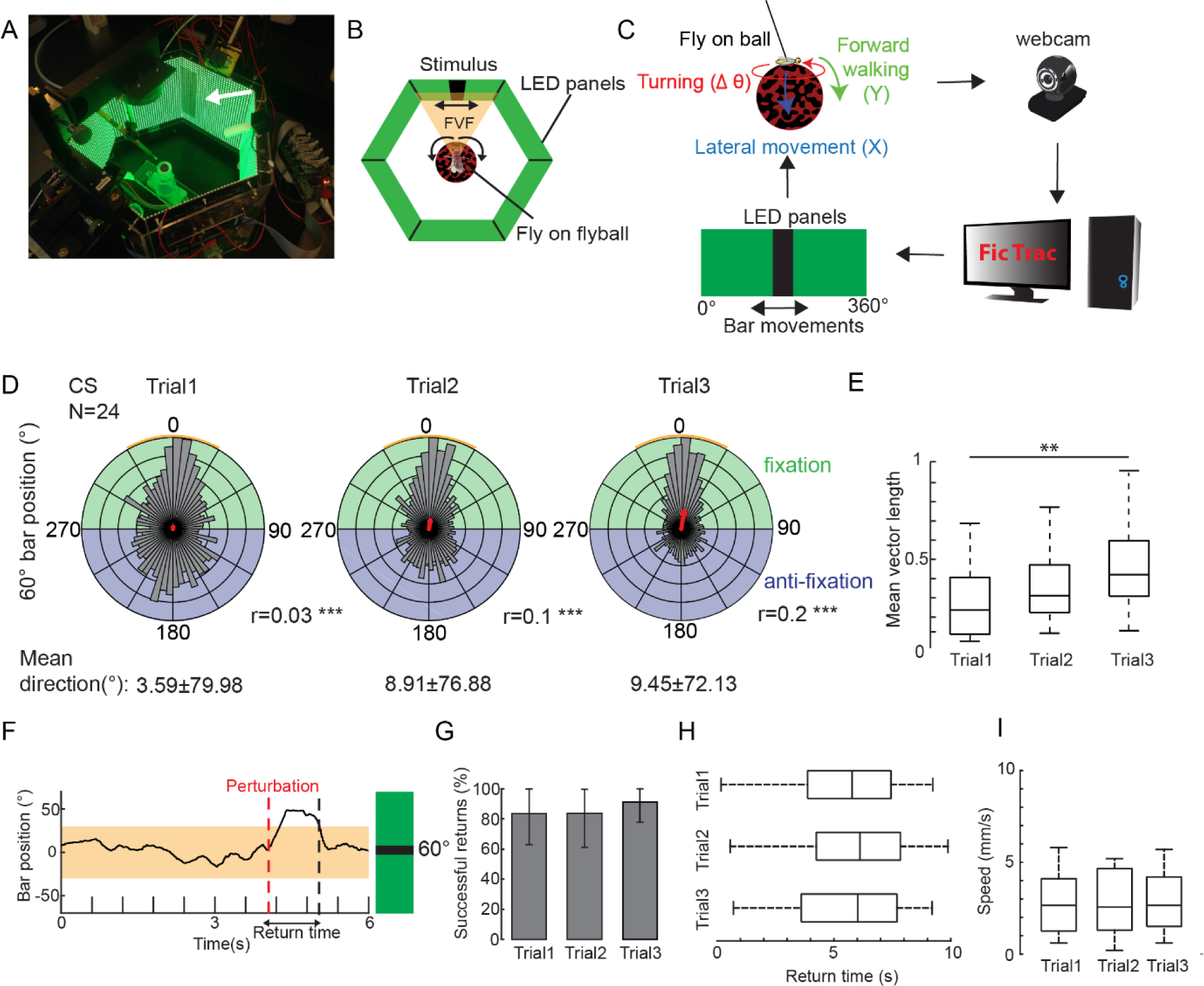
Flies fixate on a visual stimulus in a 360° LED arena. A) Virtual reality arena consisting of 6 LED panels arranged in a hexagonal shape. B) Fly is positioned on an air supported ball in the center of the LED arena. The frontal panel is defined as the frontal visual field (FVF). B) Rotations of the ball (ΔΘ) as well as forward walking (Y) and lateral movements (X) are captured via a webcam and recorded with FicTrac. A custom written Python program interacting with FicTrac translates ball movements into movements of the visual stimulus in the arena. D) Flies show increasing fixation towards a 60° bar (N=24, Rayleigh test, *p<0.05); red arrow= mean vector, r=mean vector length. Orange shade between 360° and 30° is the FVF. E) Mean vector length increases with increased trial number (N=24, Kruskal-Wallis test, ** p<0.01). F) Example of a fly reacting to a stimulus perturbation. Black trace= bar position, orange shade= FVF G) Successful returns after perturbations. (Error Bars: SD, ANOVA, α=0.05). H) Time to return stimulus in the FVF after a perturbation (successful returns only). (Kruskal-Wallis test, α=0.05). I) Average walking speed per trial (Kruskal-Wallis test, α=0.05). N=number of animals. CS=Canton(S) wildtype flies. n.s.= not significant

### Flies navigate through a virtual maze to reveal visual preferences and aversions

We next investigated visual decision-making in this paradigm. Not all visual stimuli are intrinsically attractive for *Drosophila* (Maimon, Straw and Dickinson, 2008). In order to better understand visual preferences in flies, we implemented a virtual choice maze used previously to study visual preferences in honeybees (Van De Poll *et al.*, 2015). In this previous experiment, bees were able to choose recurrently between 12 visual stimuli, which were green bars flickering at different frequencies. This operant approach revealed a clear preference/aversion profile for specific visual flickers in bees. We implemented a similar recursive approach for *Drosophila* flies, to determine visual preferences or aversions among 12 different-sized bars, presented in paired competition with one another. The bars were all dark on a lit green background, 15° wide and between 3.75° and 60° high (Fig. 2A). The flies walked on a fictive path along the edges of a (virtual) dodecahedral structure (Fig. 2B, green arrows). The faces of the structure represented the different visual objects (Fig. 2A, B, bar height in degrees). At any time, the fly was thus presented with two competing objects (faces, in Fig. 2B) 180° apart (Fig. 2C). Flies fixated on one or the other object, and after walking for 7cm, a decision was arrived by the program algorithm (see Methods) at depending on which object was most fixated upon (perturbations occurred throughout, as above, to ensure this was an active choice). The most fixated object was retained and the less fixated object was replaced by another object, represented as the next adjacent face on the dodecahedron structure (Fig. 2B). An experiment lasted until at least 80% of all stimuli were seen, or a minimum of 45 minutes per fly, yielding an average proportioned choice profile (Fig. 2D, See Methods).

**Figure 2.**
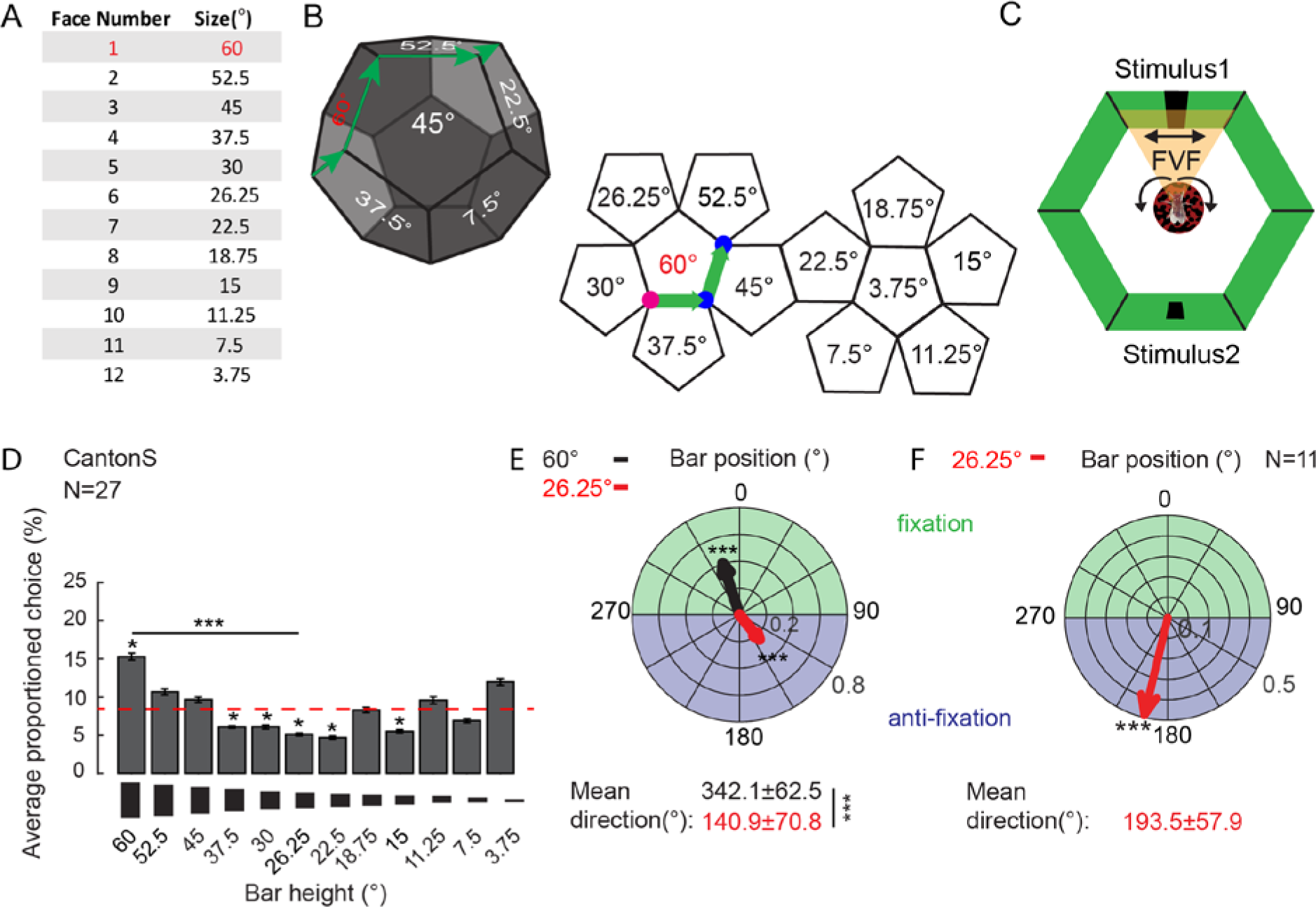
Flies show specific choice behaviour towards different sized bars. A) Each face number in the multiple-choice design is assigned to a stimulus height. B) Geometrical structure of the virtual maze. Left: Every surface of the dodecahedral structure (face) represents one distinct stimulus. Green arrow: virtual path. Faces on both sides of the path represent the displayed stimuli. Right: unwrapped virtual maze. Pink dot: starting point of experiment. Fly is first confronted with a 60° (red) and 37.5° stimulus. Blue dot: first decision point. The 37.5° bar gets replaced by the 45° bar until the next decision is made. C) In the arena the visual stimuli are locked to be 180° apart. The stimulus’ movement is locked to the fly ball’s movement. D) Average proportioned choice for all animals for all presented stimuli. Asterisks above bars represent significant differences from chance level (red dashed line, 8.3%, Wilcoxon rank-sum test, α=0.05). E) Mean direction of bar positons of 60° (black arrow) and 26.25° (red arrow) within the LED arena (Rayleigh-test for significance of mean direction, Watson-Williams test for difference between mean directions) F) Mean direction of bar position of 26.25° bar within the LED arena (Wilcoxon-rank sum test, α=0.05). N=number of animals, *p<0.05, ***p<0.001, Error bars=s.e.m.

Our closed-loop visual competition experiments revealed that wild-type, female *Drosophila* flies selected the large 60° bar above chance level (red dashed line at 8.3%, Fig. 2D). Interestingly, flies seemed to select medium-sized bars (37.5°-22.5°) significantly below chance level, suggesting these are visually repulsive to them. Calculating the mean direction vector for the stimuli revealed that the 60° bar was indeed mostly positioned in the FVF (342.1±62.5°). In contrast, medium-sized bars were positioned behind the fly: for example, the 26.25° bar was positioned in the opposite direction (140.9°±70.8°) on average (Fig. 2E). The mean direction for the largest (60°) bar and the medium (26.25°) bar were significantly different from each other (Fig. 2E), and the 60° bar was chosen significantly more often than the 26.25° bar (Fig. 2D). Having found that the 26.25° bar was consistently avoided when presented in competition with other bars, we then asked whether it was aversive even when presented on its own. Indeed, when we presented the 26.25° bar on its own (as in Fig. 1 for the 60° bar), flies displayed clear anti-fixation behavior (Fig. 2F). This confirms that the 26.25° and 60° bars are indeed visually ‘repulsive’ and ‘attractive’, respectively, and that our operant virtual maze design can effectively uncover these innate visual preferences.

We next investigated whether these innate visual preferences persisted through life, in older female flies. Older flies (17-40 day) displayed a remarkably similar choice profile as the younger (5-10 day) flies (Fig. 3A), again choosing the 60°bar significantly more often than the 26.25° bar. Interestingly, the smallest bar (3.75°) became attractive to older flies, to a similar level as the largest bar. Older flies showed a significant fixation for the 60°bar and an anti-fixation for the 26.25° bar, with a significant difference in mean vector direction between these objects (Fig. 3B). However, there was no significant difference in mean vector length between the young and old dataset, for these two visual objects (Fig. 3C). This indicates that the quality of fixation (and anti-fixation) remains robust with age. Interestingly, old age significantly decreased novelty-seeking behavior in this paradigm (Fig. 3D, E). Every visual choice represents two different historical contingencies: either a continued selection of a preferred object (a continuation choice) or a selection of a novel object that has just replaced a previously non-preferred object (a novelty choice) ((Van De Poll *et al.*, 2015) see Methods). Younger flies fixated upon the smaller 26.25° bar more often when it was novel (Fig. 3D), suggesting that novelty could override repulsion. In contrast, older flies mostly displayed continuation choices (Fig. 3E), and showed significantly less novelty-seeking behavior in general for all objects (Fig. 3F). These experiments show that innate visual preferences remain robust through a fly’s life, without deterioration in fixation behavior, although more flexibility in younger flies is evident.

**Figure 3.**
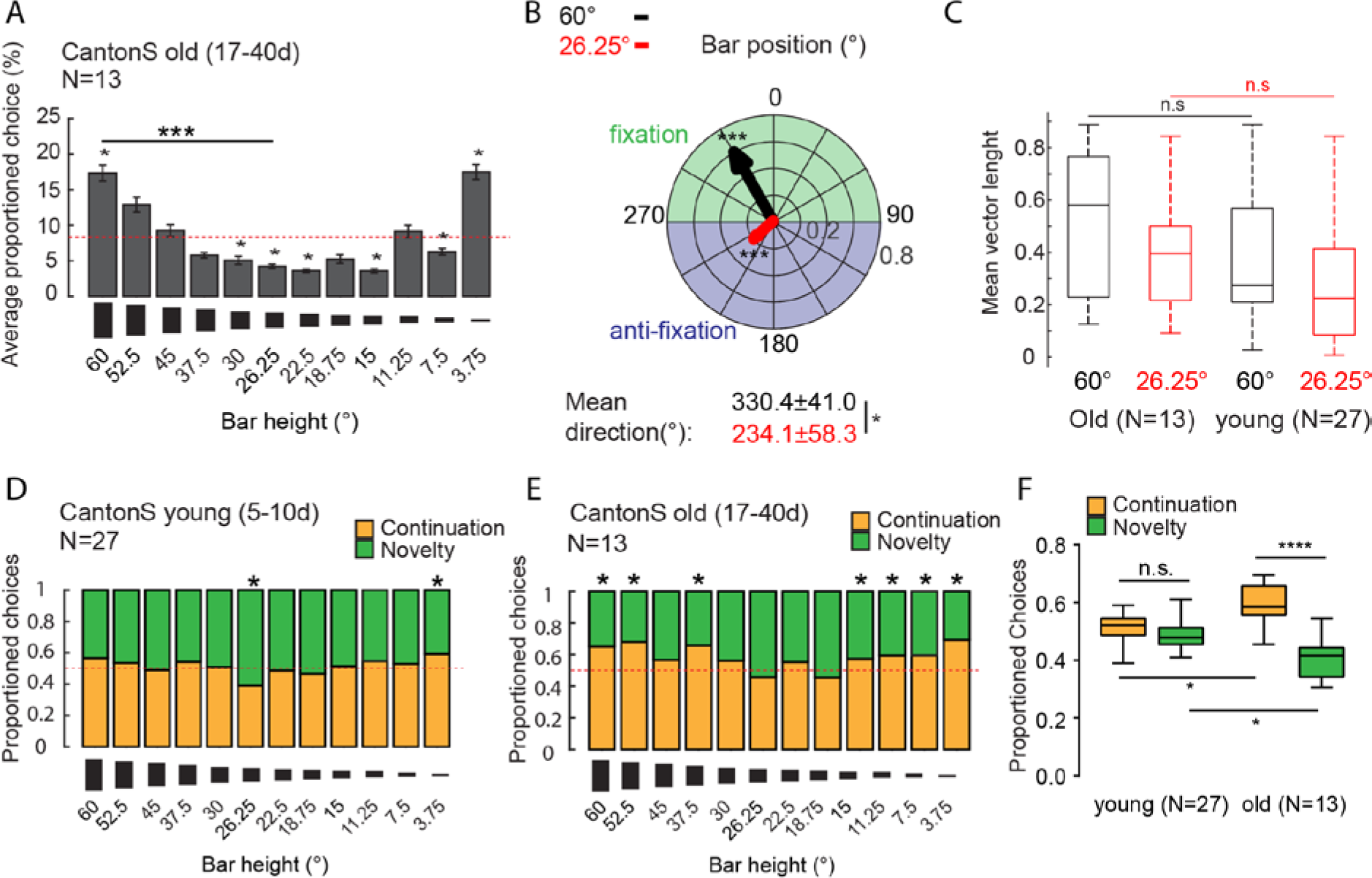
Increasing age of flies has no effect on choice behaviour. A) Choice profile of 17-40 day of (past eclosion) female CS flies. Red dashed line: 8.3% chance level. (Rank-sum test compared to chance level for single stimuli choices, Kruskal-Wallis test to compare between stimuli, α=0.05) B) Mean direction of bar positions for the 60° (black arrow, mean vector) and 26.25° (red arrow, mean vector) bars. (Rayleigh test for mean vector lengths, Wilcoxon rank-sum test for comparison of mean direction, α=0.05) C) Mean vector length for the 25.25° bar and the 60° bar for 17-40 day old flies and young 5-10d old flies (Wilcoxon rank-sum test, α=0.05). D) Proportioned novelty and continuation choices for visual stimuli of 17-40 day old flies. Red dashed line=50% chance level. E) Proportioned novelty and continuation choices for visual stimuli of young 5-10d old flies. F) Pooled novelty and continuation behaviour for old and young flies. Red dashed line = 50% chance. Wilcoxon rank-sum test, α=0.05 for nonparametric data. ANOVA for parametric data (F). N=number of animals. Error bars=s.e.m., *p>0.01, **p>0.01, ***p>0.001, n.s.= not significant

### Repulsion and attraction for different-sized objects remain robust under different color, light, and contrast conditions

A recent study investigating visual attention in *Drosophila* showed that background luminosity and color affects visual attraction and aversive behavior in flies (Koenig, Wolf and Heisenberg, 2016). To test whether this might be the case for the visual objects in our closed-loop paradigm, we ran our virtual maze experiments under different light, contrast and color conditions (Fig. 4). Changing the background color from green (RGBA: 0.0, 1.0, 0.0, 1.0) to cyan (RGBA: 0.0, 0.58, 0.58, 1.0) and maintaining the same luminosity (3279 Lux) did not alter the choice profile of young wild-type female flies, although more significant effects were noted (Fig. 4A). The 60° bar was still chosen significantly more often than the 26.25° bar, and the mean direction of each bar position for these objects was still significantly different (Fig. 4A, right box plot). Behavioral processes were also preserved under these different conditions: the 60° bar was mostly chosen as a ‘continuation’ event, whereas the 26.25° bar was fixated upon mostly if it was novel (Fig. 4B). Changing the luminosity of the cyan background (RGBA: 0.0, 0.4, 0.4, 0.8, 565 Lux) and the color of the bar from a high-contrast black to an equal-luminous red (RGB: 151, 0, 0, 550 Lux) still revealed a robust and qualitatively similar choice profile (Fig. 4C), indicating that object shape (rather than luminosity) was being selected. When we presented competing dark objects on a red background (RGBA: 0.2, 0.0, 0.0, 1.0), the choice profile became flat (Fig. 4D), showing no significant preferences for any object. *Drosophila* have little perception of red-shifted light, due to the lack of corresponding photoreceptors (Garbers and Wachtler, 2016), so it is not surprising that they could not perceive the dark objects in this context. Increasing the contrast by increasing the red background luminosity (RGBA: 0.75, 0.0, 0.0, 1.0) still showed a flat choice profile with no significant differences between bar choices as well as mean directions (Fig. S1). In addition to showing an absence of object perception in this context, these experiments indicate that the choice profiles revealed earlier are not an artifact of the maze geometry; only visible objects revealed a significant choice profile.

**Figure 4.**
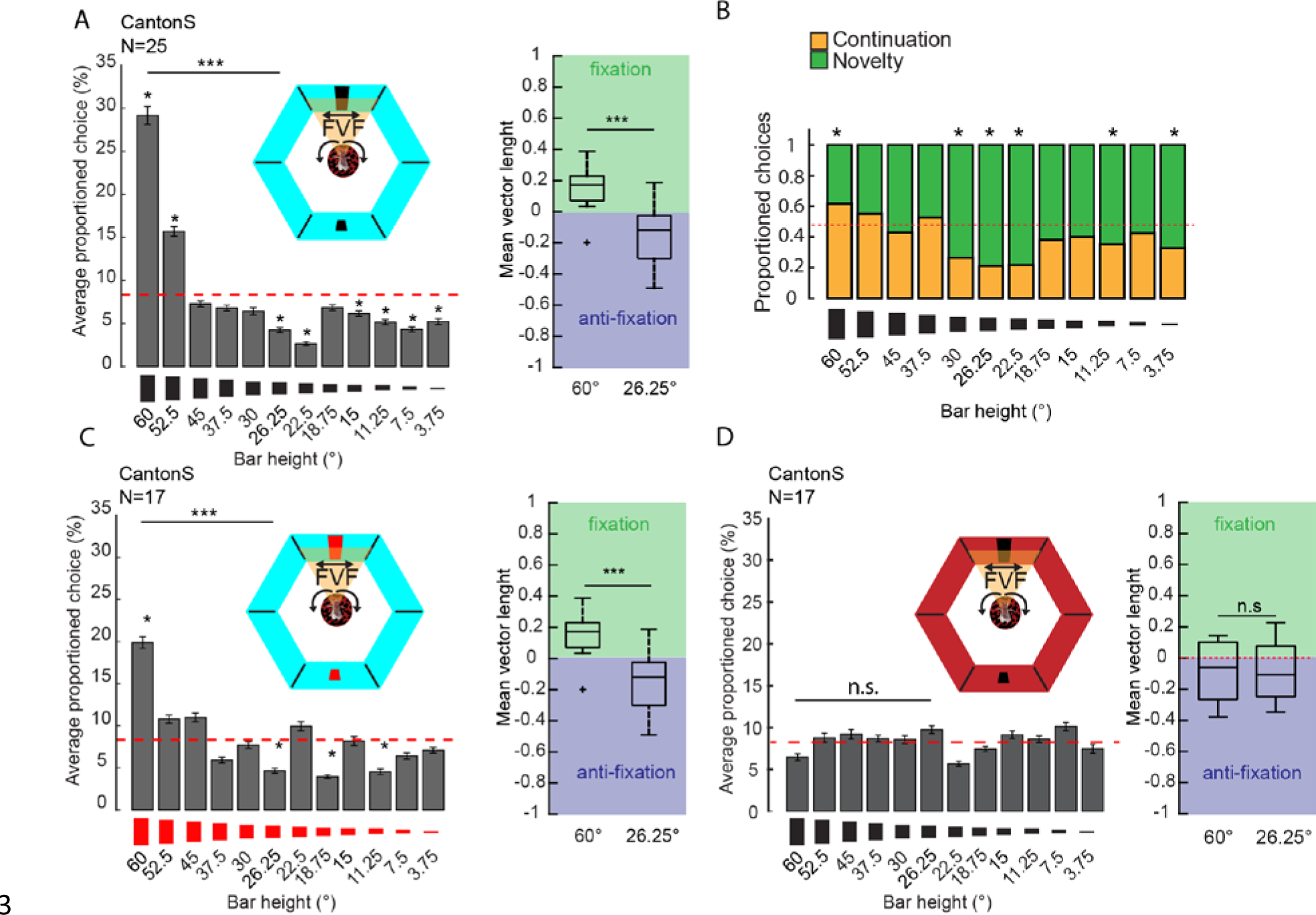
Characteristic choice behaviour for different-sized visual stimuli stays robust under different light/colour conditions. A) Left: Averaged proportioned choice for 12 different visual stimuli on cyan background (same luminosity as the green background, see Methods). Right: mean vector length of pooled 60° bar and 26.25° bar positions within the LED arena. B) Novelty and continuation choice profile for choice paradigm with cyan background. Red dashed line=50% chance level C) Average proportioned choice for red visual stimuli on cyan background (see Methods). Right: mean vector length of pooled 60° and 26.25° bar positions within the LED arena. D) Averaged proportioned choice for black visual stimuli on red (low contrast) background (see Methods). Right: mean vector length of pooled 60° and 26.25° bar positions within the LED arena. N= number of animals, red dashed line= 8.3% chance level (A,C-D), 50% chance level (B), Wilcoxon rank-sum test for comparison of mean vector length, Kruskal-Wallis test for comparison between averaged proportioned choices, Wilcoxon rank-sum test for comparison to chance level (A, C, D). α=0.05, *p<0.05, ***p<0.001.

### Pre-exposure to a repulsive object affects choice behaviors but not fixation

Habituation can have an effect on valence-based decisions (Rangel, Camerer and Montague, 2008). We therefore next investigated whether we could modulate the attractive and repulsive responses towards visual stimuli, by habituating the fly to these stimuli prior to running the virtual choice maze experiment. To test this, we pre-exposed flies to either a single 60° bar or a 26.25° bar for 3 consecutive 2min trials (Fig. 5). For the attractive stimulus (60°), we found that pre-exposure resulted in a similar choice profile (Fig. 5B, left), compared to non-habituated flies (Fig. 2). The mean direction of both the attractive and aversive bar positions within the LED arena was also unchanged: the 60° bar was fixated and the 26.25° bar was anti-fixated (Fig. 5C, left). On the other hand, pre-exposure to the repulsive 26.25° bar (Fig. 5A, right) had a greater effect on the average choice profile of the flies. Now, the 60° bar was not chosen significantly above chance level, and the 26.25°bar was also not chosen significantly below chance level as it was the case in Fig. 2 (Fig. 5B).

**Figure 5.**
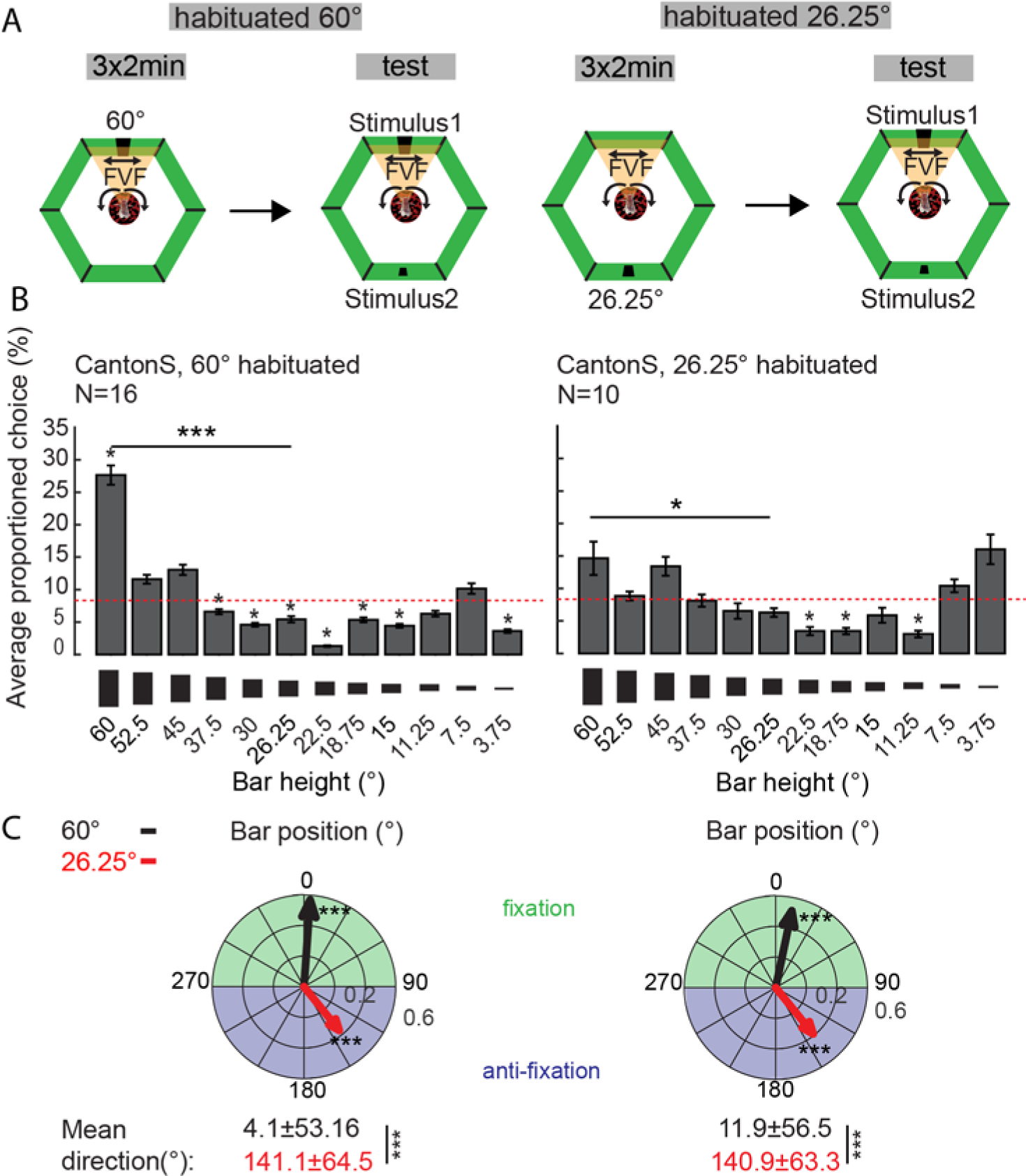
Prior exposure to an attractive and repulsive visual stimulus has different effects on choice behaviour. A) Experimental setup. Animals are exposed to a single visual stimulus for 2 minutes in three consecutive trials (left: attractive 60°, right: repulsive 26.25°). After this pre-exposure, flies were exposed to in the virtual maze, as in Fig. 2. B) Left: Averaged proportioned choice profile for visual stimuli after pre-exposure to the 60° bar Right: Averaged proportioned choice profile for visual stimuli after pre-exposure to the 26.25° bar C) Left: Mean direction and mean vector lengths of 60° (black bar) and 26.25° (red bar) after pre-exposure to 60° bar Right: Mean direction and mean vector lengths of 60° (black bar) and 26.25° (red bar) after pre-exposure to 26.25° bar N= number of animals, Wilcoxon rank-sum test, α=0.05 (B), Rayleigh test for mean vector length, and Watson-Williams test for mean direction (C), *p<0.05, ***p<0.001. Error bars =s.e.m., red dashed line=chance level 8.3%.

However, the flies still preferred the 60° bar significantly over the 26.25° bar (Fig. 5B, right). This was also reflected by the overall positions of both bars within the arena during the experiment: the 60° bar was still significantly within the FVF and the flies showed anti-fixation for the repulsive 26.25° bar (Fig. 5C, right). These results suggest that prior experience, even if relatively brief (6min), can affect valence assignations in the subsequent recursive choice paradigm. However, innate object preferences still appear quite robust.

### Activation of dNPF reduces anti-fixation towards a repulsive visual stimulus

In the preceding experiments, we have attempted to query the fly’s motivational state by examining effects of age, habituation, and stimulus parameters. We next attempted to modulate visual preferences by directly manipulating *Drosophila* brain circuits that have been associated with motivational states, such as neurons that express neuropeptide F (dNPF) (Shao *et al.*, 2017). dNPF has been associated with reward in *Drosophila* (Krashes *et al.*, 2009), by for example altering feeding behavior (Chung *et al.*, 2017), social behavior (Wu *et al.*, 2003), and olfactory learning (Krashes *et al.*, 2009; Chung *et al.*, 2017; Shao *et al.*, 2017). It is unclear however whether this neuropeptide also plays a role in gating information about the valence of visual stimuli. In the context of our recursive choice maze paradigm, we tested whether the previously-established preference profile for object size could be modulated by acute activation of dNPF neurons, by expressing the red-light shifted channelrhodopsin variant ‘CsChrimson’ (Klapoetke *et al.*, 2014) in dNPF-expressing neurons (Fig. 6A). To ensure the rewarding stimulus was only associated with one object, we transiently activated dNPF-expressing neurons only when flies fixated on the small aversive (26.25°) bar (Fig. 6B, C), in the context of our recursive choice maze paradigm. The red light was off (i.e., NPF-expressing neurons were not activated) when the fly fixated on any other visual stimulus. Control animals were not fed retinal, a food supplement required for light-induced activation of the channelrhodopsin (see Methods).

**Figure 6.**
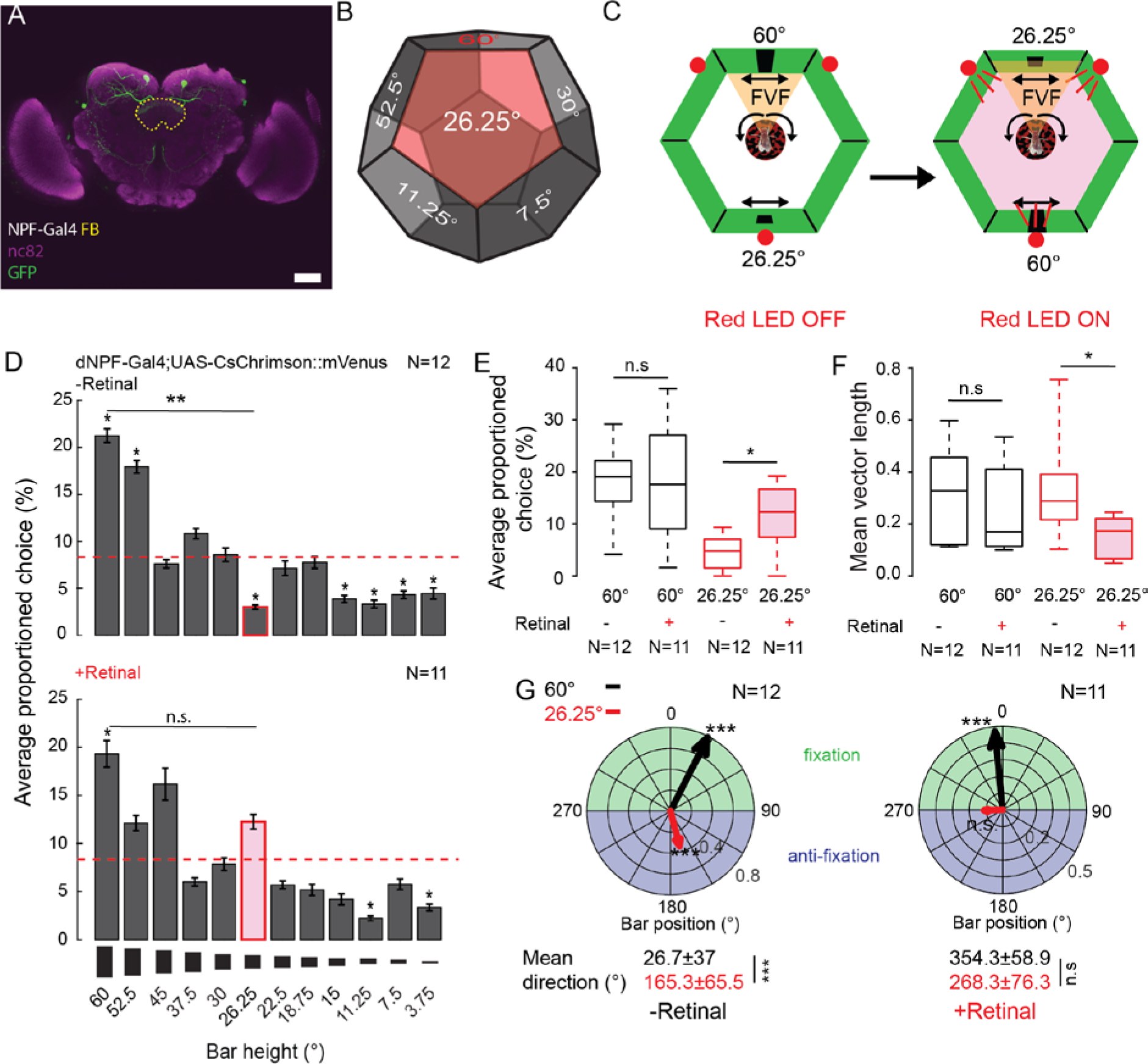
Optogenetic activation of dNPF alters choice behaviour for repulsive visual stimuli. A) dNPF-circuit (green/GFP). A subset of dNPF neurons have projections to the fan-shaped body (yellow dashed line). B) During the maze experiment only the presence of the repulsive 26.25° stimulus (red hexagonal surface) triggered the optogenetic dNPF activation by red LED lights. C) The 26.25°stimulus had to enter the FVF in order to trigger red LED activation (Red LED ON). D) Averaged proportioned choice profile of the control flies (non-retinal fed – RET) and the dNPF activated flies (retinal fed, +RET). E) Averaged proportioned choice of the 26.25° and the 60° bar with (red) and without (black) LED activation for both groups (+Retinal, -Retinal) F) Mean vector length, representing the distribution of the 60° and 26.25° bar positions in the LED arena with (red) and without (black) LED activation for both groups (+Retinal, -Retinal).(G) Left: Mean direction and vector length of the 60° (black) and 26.25° (red) bar positions within the LED arena, control group. Right: Mean direction and vector length of the 60° (black) and 26.25° (red) bar positions within the LED arena during LED activation for the 26.25° bar, retinal-fed group. N=number of animals. Error bars=s.e.m., Red dashed line=Chance level 8.3%, Wilcoxon rank-sum test, α=0.05 (D,E,F,G) Rayleigh test for mean vector length (G), Watson-Williams test for mean direction, α=0.05 (G), *p<0.05, **p<0.01,***p<0.001, n.s.=not significant.

We found that control flies displayed a similar preference profile as wild type (CS) flies (Fig. 6D, upper panel), despite the red light turning on when the 26.25° bar happened to be in the fly’s FVF. In contrast, activation of dNPF-expressing neurons in retinal-fed flies led to a loss of repulsive behavior towards the 26.25° bar, while the larger bar remained attractive (Fig. 6D, lower panel). Activation of dNPF-expressing neurons had no effect on selection of the 60° bar, but selection for the 26.25° bar was significantly increased (Fig. 6E). This was also reflected by the vector length and orientation, which remained unchanged for the large bar but decreased (due to decreased repulsion) for the smaller bar (Fig. 6F, G). Altered behavior toward the smaller bar was not due to altered walking speed, which remained unchanged (Fig. S2 2A). Increased attraction to the smaller bar was also evident in retinal-fed flies returning this object to their FVF more often, compared to controls, after a perturbation event (Fig. S2 B). Thus, negative visual valence can be eliminated by acute dNPF circuit activation.

### Activation of dNPF neurons transiently reduces aversion and is a positive cue

Since activation of the dNPF pathway is associated with olfactory learning in *Drosophila* (Krashes *et al.*, 2009), we next tested whether operant activation of the dNPF circuit in our paradigm could result in visual learning. For this purpose, we devised a closed-loop learning assay (Fig. 7A) where we rewarded the smaller (26.25°) bar in competition with an unrewarded larger (60°) bar (see Methods). We first confirmed our previous result showing that over 3 consecutive 2min trials, flies chose the large bar significantly more often than the 26.25° bar (Fig. 7B, Trial 1-3), before activating dNPF-expressing neurons. Training began in trial 4, by turning on the red light only when the small bar was in the frontal visual field.

**Figure 7.**
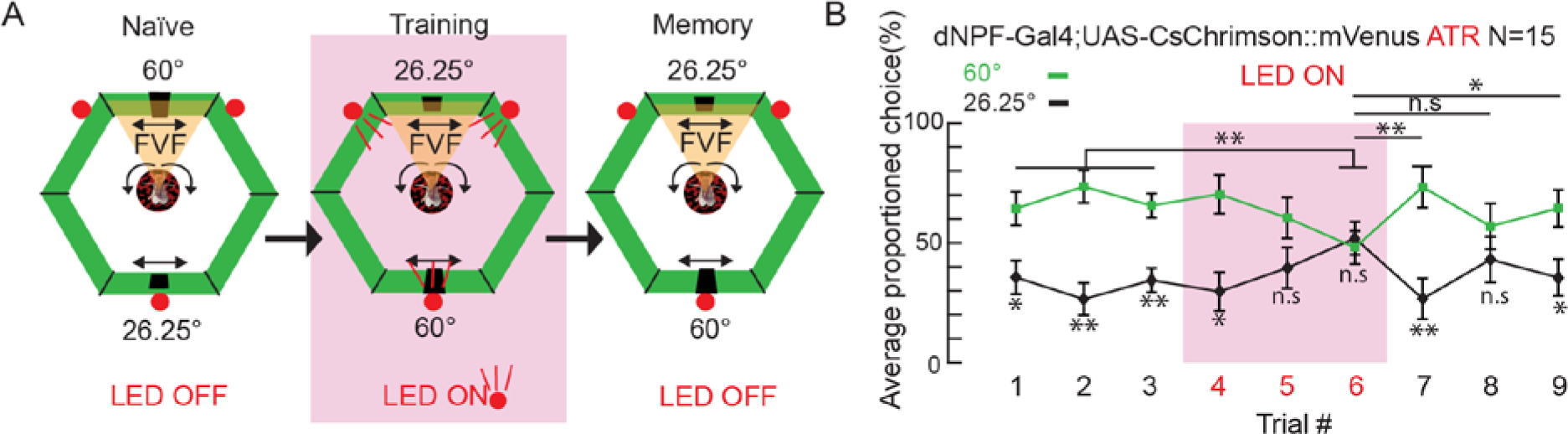
Activation of dNPF transiently reduces negative valence. A) From left to right: Naïve: retinal-fed flies were tested for their baseline fixation and anti-fixation behaviour towards the competing large 60° bar and the smaller 26.25° bar for 2 minutes in three consecutive closed-loop trials. Training: three consecutive 2 minute trials, wherein positioning of the 26.25° bar in the FVF caused the activation of red LEDs. Memory: three consecutive 2min trials, without LED activation. B) Averaged proportioned choices for 60° (green) bar and 26.25° (black) bar for all trials. Red area indicates training phase with LED activation when 26.25° bar was in the FVF. N=number of animals Wilcoxon-rank sum test between averaged proportioned bar choices, Kruskal-wallis test between trials, *, p<0.05,**, p<0.01, *** p<0.001, n.s.= not significant. Error bars= s.e.m.

Flies lost their aversive behavior towards the small bar by trial 5 (Fig. 4B), showing no significant difference in mean choice behavior in trials 5 and 6. However, as soon as the dNPF circuit was no longer being activated (trial 7), flies returned to their innate preferences. This suggests that the 26.25° bar was rendered transiently attractive by the operant reward paradigm – and was at least equivalent in valence to the 60° bar – but that the valence effect did not persist beyond the training session.

To ensure that we had altered valence rather than merely causing indifference to the competing bars, we devised a different operant learning experiment, where flies were rewarded (by activating NPF-expressing neurons) only when they placed an object (the 60° bar) in a specified quadrant of the arena (Fig. 8A). Flies were first tested for baseline fixation behavior, which for the larger bar is naturally directed towards the FVF (Fig. 8A, B, first panel). To test for operant learning, dNPF-expressing neurons were activated only when the bar was positioned to the right of the fly (between 60° and 120°, Fig. 8A, second panel). Accordingly, flies kept the bar significantly more often on their right side (Fig. 8B, second panel). We confirmed this result by rewarding bars positioned behind the fly or to the left, with flies accordingly placing the bar in those respective quadrants in closed loop (Fig. 8 A, B third and fourth panels). This shows that activation of the dNPF circuit is indeed rewarding, and not simply abolishing visual behavior. Additionally, after every experiment, we asked whether flies continued to place the bars in these previously-rewarded locations, by testing the flies again in a two-minute trial without activating dNPF neurons (see Methods). However, flies immediately returned to keeping the stimulus in their FVF after the operant training (Fig. 8 A, B, 1. TRIAL LED OFF). We conclude that in this paradigm, activation of the dNPF circuit provides a transient positive cue that does not have a lasting effect on visual learning.

**Figure 8.**
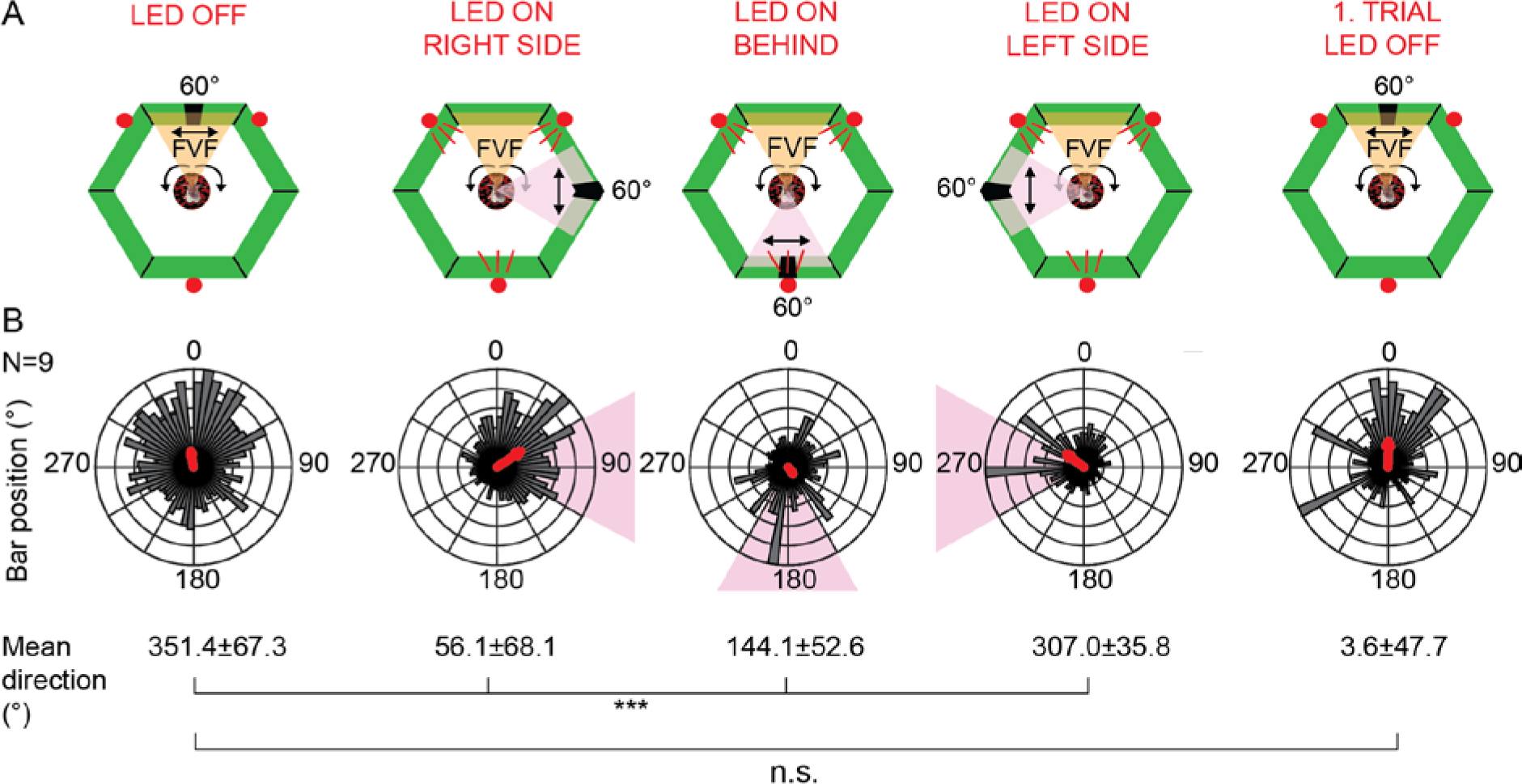
Acute activation of the dNPF circuit is a positive cue. A) Experimental setup. Red LEDs were activated when the bar was on the left, back and right side (red shade). Orange area indicated the FVF B) Mean directions and mean vectors (red arrow) of bar positions during the experiment. Each polar plot is the average of 3 consecutive 2 minute trials for all animals. 1 Trial LED OFF is the average of 3 2 minute trials for all animals. Red area indicates where the bar needed to be to trigger the activation of the red LEDs. Darker histograms in polar plots represent binned bar distributions. 1.Trial LED OFF represents the first 20 seconds after the optogenetic activation. N=number of animals, *** p<0.001, n.s. = not significant, Watson-Williams test.

## Discussion

All animals display strong innate preferences, being attracted to some stimuli and repulsed by others, which influences their ultimate decisions and actions. With odors, this easily relates to chemicals relevant to an animal’s survival in a specific environment: the smell of rotten food is repulsive to humans but attractive to a fly. Visual stimuli are more difficult to assign valence, as these tend to be highly context dependent (Heisenberg, Wolf and Brembs, 2001; Brembs and Wiener, 2006). Simple visual parameters, such as responses to light intensity (Menzel, 1979; Reichert and Bicker, 1979) and color (Menne and Spatz, 1977; Morante and Desplan, 2008), have been well studied in *Drosophila melanogaster*. In contrast, fly responses to visual objects with multiple features are less well understood (Paulk, Millard and van Swinderen, 2013). While there has been a considerable amount of research done on visual learning in tethered, closed-loop flight paradigms, these experiments are typically not concerned with uncovering innate preferences, but rather focus on feature discrimination of visual objects of a possible equivalent valence such as upright ‘T’s and upside down ‘T’s (Heisenberg, Wolf and Brembs, 2001; Liu *et al.*, 2006; Paulk, Millard and van Swinderen, 2013). Indeed, most visual learning paradigms in animals are agnostic of the larger valence landscape wherein experimental stimuli reside – or even whether they are attractive, aversive, or neutral – but rather settle on robust responses that produce reliable behavioral readouts. For visual decision-making however, some knowledge about innate valence is important for better understanding responses to different stimuli (Guitart-Masip *et al.*, 2014).

In this study, we use a closed-loop visual paradigm for walking flies to show that flies find large bars innately attractive and smaller bars repulsive. This confirms previous work done in closed-loop flight (Maimon, Straw and Dickinson, 2008), indicating surprisingly entrenched valence effects for these simple visual objects. We used a virtual reality maze paradigm, previously developed for walking honeybees (Van De Poll *et al.*, 2015), to place these preferences within a larger valence spectrum for object size, particularly bars of different heights but same width. In this paradigm, flies are able to reveal their visual preferences, by iteratively selecting competing objects in a recurrent binary choice design. We find that visual preference profiles are remarkably robust, with larger bars remaining more attractive and smaller bars repulsive even as flies age, or when they are exposed to different visual experiences, such as different background colors, and luminosities. Earlier studies have shown that these visual stimulus parameters can affect a fly’s attention and therefore learning behavior (Koenig, Wolf and Heisenberg, 2016). We did find, however, that flies fixate on innately repulsive objects if they are perceived as novel, suggesting that an internal switch exists for over-riding these deeply-entrenched valence effects. This novelty effect that was also observed in honeybees for innately aversive visual flickers (Van De Poll *et al.*, 2015).

An interesting observation was made in older flies compared to younger flies. Whereas older flies displayed similar valence profiles as younger flies, and their fixation behavior was just as robust, they were less responsive to visual novelty. This conservative behaviour in older flies has some parallels with human behaviour: aging in humans affects decision making, resulting in less impulsive and delayed choice behavior, compared to younger participants (Eppinger, Hämmerer and Li, 2011; Eppinger, Nystrom and Cohen, 2012). Such superficial similarities in valence-based decision making between older flies and older humans might not be surprising considering the likelihood of homologous systems being involved in decision-making in both species (Bogacz and Gurney, 2007; Strausfeld and Hirth, 2013; Barron *et al.*, 2015; Barron and Klein, 2016).

One important aspect of valence-based decision making is that it is not necessarily hard-wired; it can be influenced by experience (Dickinson and Balleine, 2002; Dayan and Abbott, 2003; Rangel, Camerer and Montague, 2008). This was observed to some extent in our experiments. A relatively short exposure to a repulsive stimulus altered valence effects in the following ˜1hr in the virtual choice maze. This resulted in a more blunted valence profile, compared to the profile following pre-exposure to an attractive stimulus. This suggests that even a brief experience can alter decision-making over a long period of time. Typically, outside of virtual reality environments, it is impossible to ascertain an animal’s exact previous experience.

We found that a reward circuit in the fly brain might play a key role in governing visual decision-making. NPY is a highly conserved neuropeptide (C Wahlestedt and Reis, 1993; Feng *et al.*, 2003) that regulates motivational states in animals (Bannon et al., 2000), and the fly homolog dNPF (Garczynski *et al.*, 2002; Shao *et al.*, 2017) seems to play a similar role. In mammals, there is also evidence that NPY plays a role in suppressing anxiety and fear (Thorsell, 2000; Redrobe, Dumont and Quirion, 2002; Primeaux *et al.*, 2005; Fendt *et al.*, 2009), as well as regulating responsiveness to aversive or stressful stimuli (Bannon *et al.*, 2000; El Bahh *et al.*, 2001). In flies, dNPF has been linked to olfactory learning by regulating dopaminergic input to the mushroom bodies (Krashes *et al.*, 2009), with data suggesting that it provides a rewarding cue (Rohwedder *et al.*, 2015; Shao *et al.*, 2017). The role of dNPF input to visual centers such as the CC is less clear, although recent studies similarly propose a reinforcing neuromodulatory role (Krashes *et al.*, 2009; Chung *et al.*, 2017; Shao *et al.*, 2017). We found that optogenetic activation of dNPF-expressing neurons could indeed be used as positive reinforcement, which agrees with a recently published study (Shao *et al.*, 2017): flies could be induced to ‘place’ a visual object at different positions in the arena, by only activating the NPF-expressing neurons when flies kept the object in that specified location. Flies innately fixate on objects in their frontal visual field (Heisenberg, Wolf and Brembs, 2001; Guo *et al.*, 2015), so their capacity to also fixate to the sides suggest an attention-like effect (Sareen, Wolf and Heisenberg, 2011; Sun *et al.*, 2017) linked to reward. Consistent with this result, rewarding a repulsive object (the smaller bar) made it more attractive. Our operant conditioning experiment rewarding the repulsive 26.25° bar in competition with the 60°bar further confirmed a role for dNPF in visual decision-making in *Drosophila*, although the transient nature of this effect suggests that other systems might need to be recruited to effectively transform these altered preferences into a more persistent memory.

## Materials and Methods

### Experimental animals

*Drosophila melanogaster* were reared using standardized fly media and kept under a 12 hour light and dark cycle at 25°C. Canton-S (CS) flies were used as wild-type (control) flies. For optogenetic control of the dNPF circuit we made use of the Gal4/UAS system (Brand and Perrimon, 1993) to express red-shifted channelrhodopsin ‘Chrimson’, a non-selective ion channel, in dNPF neurons. Exposure to red light results in an activation of these ion channels and therefore an activation of the dNPF neurons. dNPF-Gal4 flies (kindly provided by Ulrike Heberlein, Janelia Research Campus, USA) were crossed with UAS-CsChrimson::mVenus(attp40) flies (kindly provided by Vivek Jarayaman, Janelia Research Campus, USA) to provide female transgenic flies used in this study. For optogenetic activation of the dNPF-neurons, Gal4;UAS-CsChrimson::mVenus flies were fed with blue-dyed 0.2mM all-*trans*-retinal (Sigma-Aldrich, St. Louis, MO, USA) food for 2 days before the experiments and kept in darkness until testing (Klapoetke *et al.*, 2014). Non-retinal fed animals were used as controls.

We used 3-11 days or 17-40 days (post eclosion) female flies. Each fly was immobilized under cold anesthesia (0.5°C) for 60s and positioned for tethering on a custom made preparation block. The flies were then glued dorsally to a tungsten rod by means of dental cement (Coltene Whaledent Synergy D6 Flow A3.5/B3) (Fig. 1B) and cured with blue light (Radii Plus, Henry Schein Dental). In order to avoid wing movements and to encourage walking of the fly, its wings were tethered to the tungsten rod by folding them against the rod and using dental cement for fixation. Additionally, to stabilize the head, it was fixed by applying dental cement to the neck of the fly (Paulk *et al.*, 2015). After tethering, the animals were provided with water and allowed to rest for about 60 minutes before testing.

### Experimental setup

The virtual reality arena was set up as described in Van de Poll *et al*. (Van De Poll *et al.*, 2015). The hexagon-shaped arena consists of six 32×32 pixel LED panels (Schenzhen Sinorad Medical Electronics Inc.) (Fig. 1A). In its center, a visually patterned, air-supported Styrofoam ball (40mg, 15mm diameter; Spotlight Ltd. Pty.) was used as walking medium for the tethered flies (Fig.1 B). For positioning of the flies on the ball, a 6-axis micromanipulator (Edmund Optics) was used. The setup was additionally illuminated by three 40W bulbs in order to provide adequate lighting for tracking the fly and ball movements by a camera (Point Grey Laboratories) at 60fps, mounted at the front of the arena. The video was further analyzed by FicTrac (Moore *et al.*, 2014), a custom made tracking software, operating in Ubuntu Linux (12.10) running on Windows 7 (SP1). To create a closed-loop environment where the fly could control the position of the stimulus, the movements of the stimuli were linked to the movements of the ball. This was achieved by linking the output (movement of the fly on the ball) of FicTrac with custom written Python (2.5) scripts (modified after Van de Poll et al. 2014) which then in turn generated the visual output with the corresponding stimulus position through VisionEgg software (Straw, 2008). FicTrac extracted the lateral movement (X), the forward movement (Y) and the rotation of the ball (turning ΔѲ) (Fig.1C) and calculated a fictive path of the fly movements which then resulted in a 1:1 translation between the movement of the ball and the rotation on the stimulus within the 360° arena (25ms delay).

To induce Chrimson activation, three orange-red LED lights (Luxeon Rebel, 617nm, 700mA, LXM2-PH01-00700) were mounted around the arena, focusing on the center of the arena. The activation and inactivation of the red LED lights was linked to the position (Fig. 6-7) and size (Fig. 6) of the stimulus, provided by FicTrac and controlled by a BlinkStick (Agile Innovative Ltd), and a LED controller board, driven by a custom written Python (2.7) script. For these experiments, position thresholds triggered the activation and inactivation of the LEDs. In Fig. 6 for example, LED activation was induced via BlinkStick when the 26.25° bar was positioned by the fly in-between 330°and 30° (FVF).

A lux meter (LX101BS) was used to determine luminosity of the background and the visual stimuli and the mean calculated from four independent measurements. The red color of the stimulus was within 1.5% error of the mean cyan background in Fig. 4. Colors of the arena were set in the custom written Python (2.7) script as RGBA values. Colors of the visual stimuli were set as RGB values in the same script.

### Behavior

All behavioral experiments were performed in closed-loop. Flies were positioned on the air-supported ball in the LED arena (Fig. 1A) and allowed to habituate to the new environment by for about 2-3 minutes.

#### Single bar fixation

In order to examine general fixation, the flies were exposed three times 2 minutes in succession to a solid black (unlit) bar on green (555nm) background. The bar was 15° (8px) wide and 60° (32px) high (Fig. 1C) or 15° (8px) wide and 26.25° (14px) high (Fig. 1A-C). If a fly was fixating on the bar, it kept the stimulus within its frontal visual field (FVF), which was defined as the width of the frontal panel (32px (60°)) (Fig. 1). Random perturbations were used in order to determine the quality of fixation. The bar was displaced every 10-30s by 60° (32px) to the left or to the right. The threshold for a successful repositioning was 10 seconds, or less.

#### Multiple choice maze

For the multiple choice maze, a set of 12 different visual stimuli was presented to the fly (Van De Poll, Zajaczkowski, Taylor, Srinivasan, & van Swinderen, 2015). The stimuli were all solid black (red) bars on green (red/cyan) background and 15° wide with different heights (Fig. 2A, B). The center of the bars was linked to the vertical center of the LED panels, so differences in bar height resulted in a symmetrical change. The flies were exposed to two competing stimuli, stimulus 1 and stimulus 2, always 180° apart, such that only one could be fixated upon at any point in time (Fig. 2C). A stimulus was regarded as successfully chosen when the fly walked a distance of 7cm and mostly fixated on one of the competing stimuli (usually 20-50s, for further details see Van de Poll et al., 2015). The unfixated stimulus was then replaced by a new stimulus (Fig. 2B). Subsequently, the fly had to choose again between the previously fixated/chosen stimulus (continuation) and the new (novelty) stimulus (Fig. 3C). This allowed us to study choice behavior in a historical context. The experiment wa s ended after the flies were exposed to at least 80% of the possible choices, which resulted in experiments between 40-60 mins typically depending on the walking speed of the flies.

#### Habituation experiment

For the habituation experiments in Fig. 4, flies were pre-exposed either to a single 60° or a 26.25° high bar for three consecutive 2 minute trials as described for the single-bar fixation experiments. Directly after the three consecutive trials the flies were presented with the multiple-choice maze as described above. There was a 10 second break between all trials where the flies walked in the arena without visual stimulation.

#### Multiple-choice maze: rewarding the 26.25° bar

For the optogenetic experiment in Fig. 5 we used the same multiple choice paradigm as described for Fig. 2. In order to activate the red LEDs the fly had to position the 26.25° bar in the FVF (330°-30°). As soon as the 26.25° bar left this area the LEDs were turned off via the BlinkStick (Agile Innovative Ltd).

#### Binary-choice paradigm: rewarding the 26.25° bar

For the learning experiments in Fig. 7 flies were presented with two competing solid black bars (60° and 26.25°) on a green background. At any time, the bars were 180° apart. Flies could fixate on one or the other. A choice was made when flies kept one of the bars for most of the time within a time frame of about 20s in their FVF. Random perturbations ensured active fixation. The experiment was divided into three parts: control, training, memory. In the control trials the flies had to choose between the two bars for 2 minutes in three consecutive trials. In the training trials (3×2min), red LEDs were activated every time the 26.25° bar was in the FVF using the BlinkStick controller board (Agile Innovative Ltd). In the last three trials (2min each) memory was tested, without the activation of the red LEDs.

#### Single-bar fixation: rewarding position and the 60° bar

For the single bar fixation experiments in Fig. 8, red LEDs were activated when the dark solid bar was at different positions in the arena (Right (60°-120°), back (150°-210°), left (240°-300°)). As in the last 2 experiments LED activation was controller by BlinkStick (Agile Innovative Ltd). Each position was tested for three consecutive 2 minute trials. Fixation was averaged across these three trials (no significant difference was found between trials, Watson-Williams-test, α=0.05, data not shown). Every time after one side was tested for fixation, we returned to a 2 minute trial of single bar fixation without optogenetic activation in order to test if there was a visible learning effect. To test for an immediate learning effect we extracted 20 seconds from this trial and analyzed the mean direction of the bar position after dNPF stimulation. After we did not observe any significant learning effect for all directions compared to control trials (fixation without LEDs activation) (Watson-Williams-test, α=0.05, data not shown) we averaged the first trial without dNPF activation after the dNPF activation for all sides and titled it 1. TRIAL LED OFF.

### Immunohistochemistry and Imaging

dNPF-Gal4,UAS-mCD8::GFP flies were collected under CO2 anaesthesia and dissected on cold 1×PBS. Samples were then fixed with 4% paraformaldehyde diluted in PBS-T (1×PBS, 0.2 Triton-X 100) for 20 min, followed by 3×20 min. washes in PBS-T. They were then blocked with 10% Goat serum (Sigma Aldrich, St. Louis, MO, USA) for 1h and incubated in primary antibody overnight. We used mouse antibody to nc82 (1:100, Developmental Studies Hybridoma Bank (DSHB) and rabbit antibody to GFP (1:1000, Invitrogen). After 3x 20 min. washes with PBS-T, secondary antibody was added and the tube was covered with aluminum foil for overnight incubation. We used AlexaFluor-488 goat anti-rabbit (1:250, Invitrogen) and AlexaFluor-647 goat anti-mouse (1:250, Invitrogen) as secondary antibodies. Following 3 final washes with PBS-T, samples were transferred to microscope slides and mounted on a drop of vectashield (Vector Laboratories, Burlingame, CA) for imaging.

Imaging was done by using a spinning-disk confocal system (Marianas; 3I, Inc.) consisting of an Axio Observer Z1 (Carl Zeiss) equipped with a CSU-W1 spinning-disk head (Yokogawa Corporation of America), ORCA-Flash4.0 v2 sCMOS camera (Hamamatsu Photonics), 20x 0.8 NA PlanApo objective was used and Image acquisition was performed using SlideBook 6.0 (3I, Inc).

### Statistics

FicTrac datasets were imported for offline analysis in MATLAB 2015b as well as in GraphPad Prism 7.0. In order to analyze fixation, bar positions were converted into polar coordinates and their mean vector length calculated using the Circular Statistics Toolbox for MATLAB (Berens, 2009). Furthermore, the distribution of the bar positions and the mean-vector length was tested for non-uniformity using the Rayleigh test (p<0.05 (*), p<0.01(**), p<0.001 (***)). Mean-vectors of fixation were weighted amongst animals and trials and tested for statistical significance using the Kruskal-Wallis test (significance level α=0.05). The mean-direction of the mean vectors was compared using the Watson-Williams (Watson and Williams, 1956; Berens, 2009). The walking speed was extracted from the XY coordinates derived from FicTrac (Moore *et al.*, 2014) and also tested for significance using the Kruskal-Wallis test with the same significance levels after weighting the datasets. Preferences for bar sizes were analyzed as proportions of the overall binary choice-experiment and their distribution was compared using a Kruskal-Wallis test with a correction by the Dunn’s multi-comparison test. These proportions were each also compared to expected chance level (8.33%) with a Wilcoxon rank-sum test. To investigate the effect of choice behavior over time, continuation and novelty choices were plotted for each animal, independent of the stimulus. Additionally, their choice behavior was weighted and proportioned for all animals (novelty *vs.* continuation) for each stimulus. Significance was calculated using a Wilcoxon-rank sum test, α=0.05, and compared between novelty and continuation. All bar plots display medians with standard errors of the mean (s.e.m.). All data was tested for normal distribution using the D’agostino-Pearson omnibus normality test. Data that passed the test for normal distribution was analyzed using a t-test and multiple datasets were compared using a one-way ANOVA with a Tukey correction for multiple comparisons.

## Acknowledgments

We thank Leonie Kirszenblat, Lucy Heap, Chelsie Rohrsheib, Adam Hines, Kai Feng, Lisa Wittenhagen and Eva Maria Reuter for helpful comments on the manuscript. We thank James Reuben for helping with behavioral experiments. We thank the QBI microscopy facility for help with imaging. This work was supported by an Australian Research Council Discovery Project grant DP140103184 to BvS and by the German Research Foundation (DFG) Research Fellowship GR 5030/1-1 to MJG.

## Author Contributions

M.J.G, J.S, and J.A. performed behavioral experiments. D.E. performed immunolabeling and imaging. M.J.G, J.A, J.S, and M.V.D.P. designed the visual paradigms and analyzed behavioral data. M.J.G and B.v.S. designed the study and wrote the manuscript.

## Data availability

All data and code used for this study are available upon request.

---

**Supplementary Figure 1.**
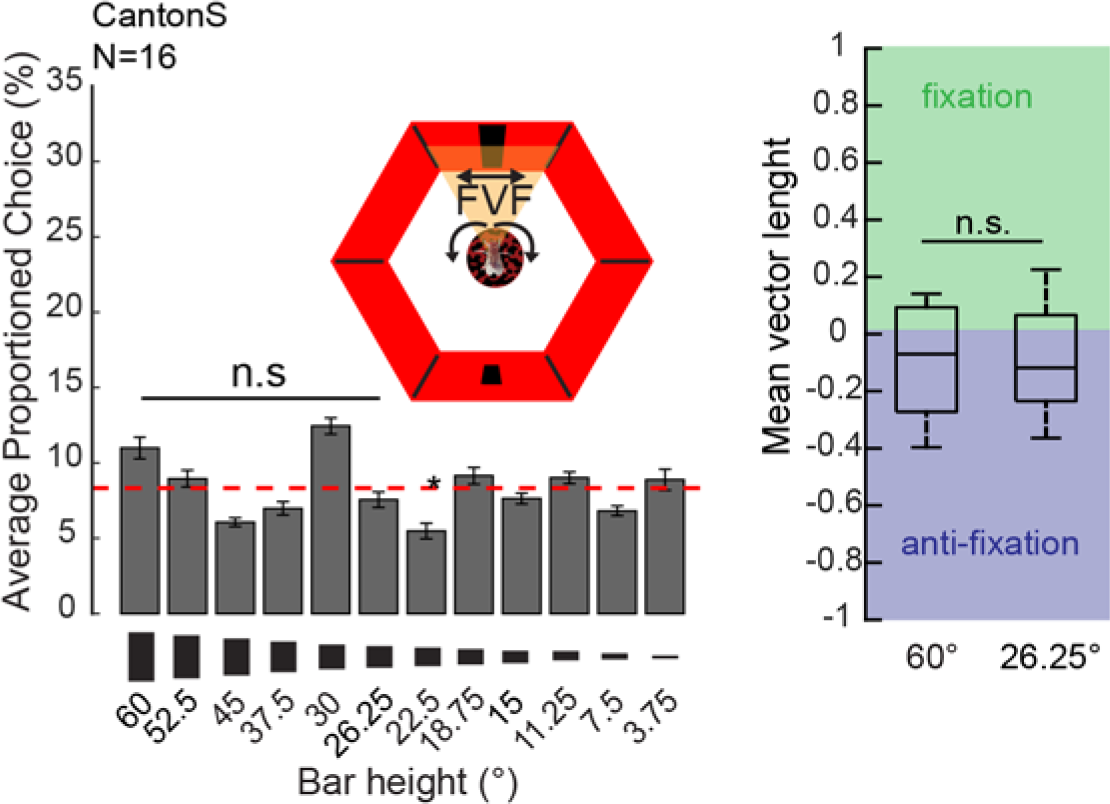
Averaged proportioned choice profile for dark stimuli on bright red background (high contrast, see Methods). Right: The mean vector length and direction for the 60° and the 26.25° bar positions within the LED arena during the experiment. Red dashed line= chance level (8.3%). For all statistics: Wilcoxon rank-sum test, α=0.05, for comparison to chance level. Kruskal-wallis test for comparison between stimuli choices, α=0.05. N= number of animals. Error bars=s.e.m.

**Supplementary Figure 2.**
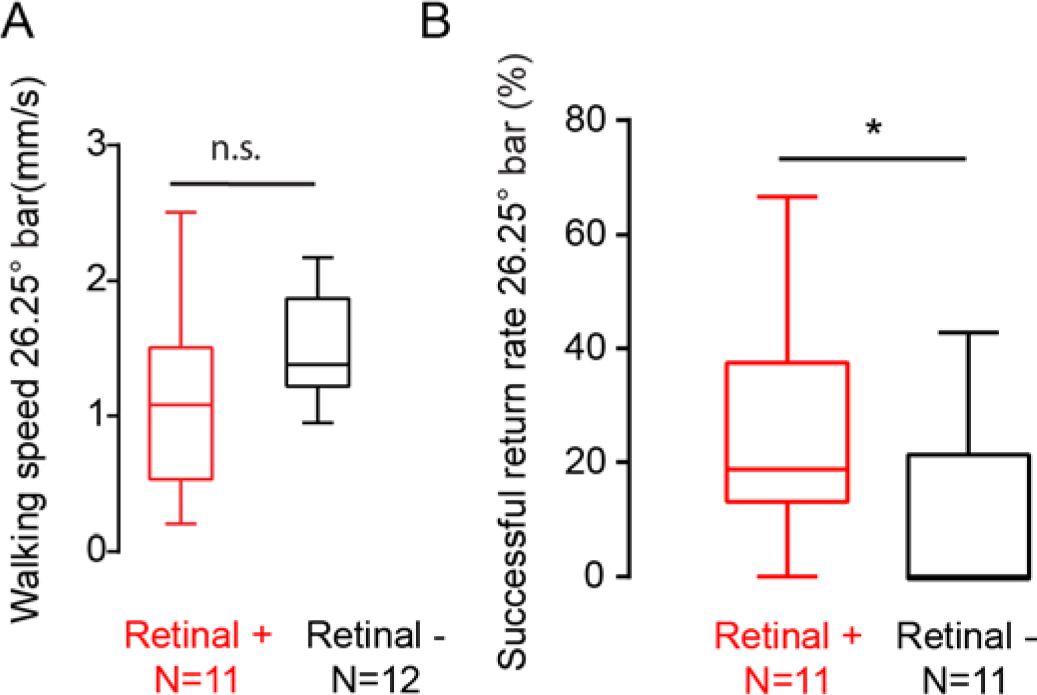
A) Averaged walking speed towards the 26.25°bar between the control group (-Retinal) and the dNPF activation group (+Retinal). t-test, p=0.6433, n.s.=not significant. B) Percentage of successful returns for the 26.25° bar for retinal fed group (Retinal +, red) and control group (Retinal -, black). *p>0.05, Mann-Whitney test, N= animals.

